# Cell-cycle status of male and female gametes during Arabidopsis reproduction

**DOI:** 10.1101/2023.02.22.529524

**Authors:** Yoav Voichek, Bohdana Hurieva, Caroline Michaud, Anna Schmücker, Zaida Vergara, Bénédicte Desvoyes, Crisanto Gutierrez, Viktoria Nizhynska, Benjamin Jaegle, Michael Borg, Frédéric Berger, Magnus Nordborg, Mathieu Ingouff

## Abstract

Fertilization in *Arabidopsis thaliana* is a highly coordinated process that begins with a pollen tube delivering the two sperm cells into the embryo sac. Each sperm cell can then fertilize either the egg or the central cell to initiate embryo or endosperm development, respectively. The success of this double fertilization process requires a tight cell cycle synchrony between the male and female gametes to allow karyogamy (nuclei fusion). However, the cell cycle status of the male and female gametes during fertilization still remains elusive as DNA quantification and DNA replication assays have given conflicting results^1–4^. Here, to reconcile these results, we quantified the DNA replication state by DNA sequencing and performed microscopic analyses of fluorescent markers covering all the phases of the cell cycle. We show that male and female gametes in Arabidopsis are both arrested prior to DNA replication at maturity and initiate their DNA replication only during fertilization.

---

As a first step, we adapted a DNA sequencing (DNA-Seq) approach^5^ that can detect even subtle DNA replication signals (Fig. 1a). To demonstrate the ability of the DNA-Seq approach in detecting DNA replication in Arabidopsis, we performed a control experiment where suspension cells were synchronized to enrich for G1- or S-phase (Extended Data Fig. 1). Genome-wide coverage in S-vs. G1-phase cells showed an increase along chromosome arms, as expected from early replicating regions (Fig. 1b, gray curve). Next, we applied our DNA-Seq approach to sperm and vegetative nuclei isolated from mature pollen grains using fluorescence-activated cell sorting (FACS)^6^. We first compared the genome-wide coverage of purified sperm nuclei with that of vegetative nuclei or root nuclei. In contrast to our control experiments, the coverage was uniform along the chromosomes (Fig. 1b, green curve, Extended Data Fig. 2). Although the genome-wide coverage pattern showed no evidence of DNA replication, a subpopulation of sperm cells may still be replicating their DNA. This would be evident as a weaker signal of DNA replication when compared with previously published replication data from Arabidopsis^7^.

**Fig. 1.**
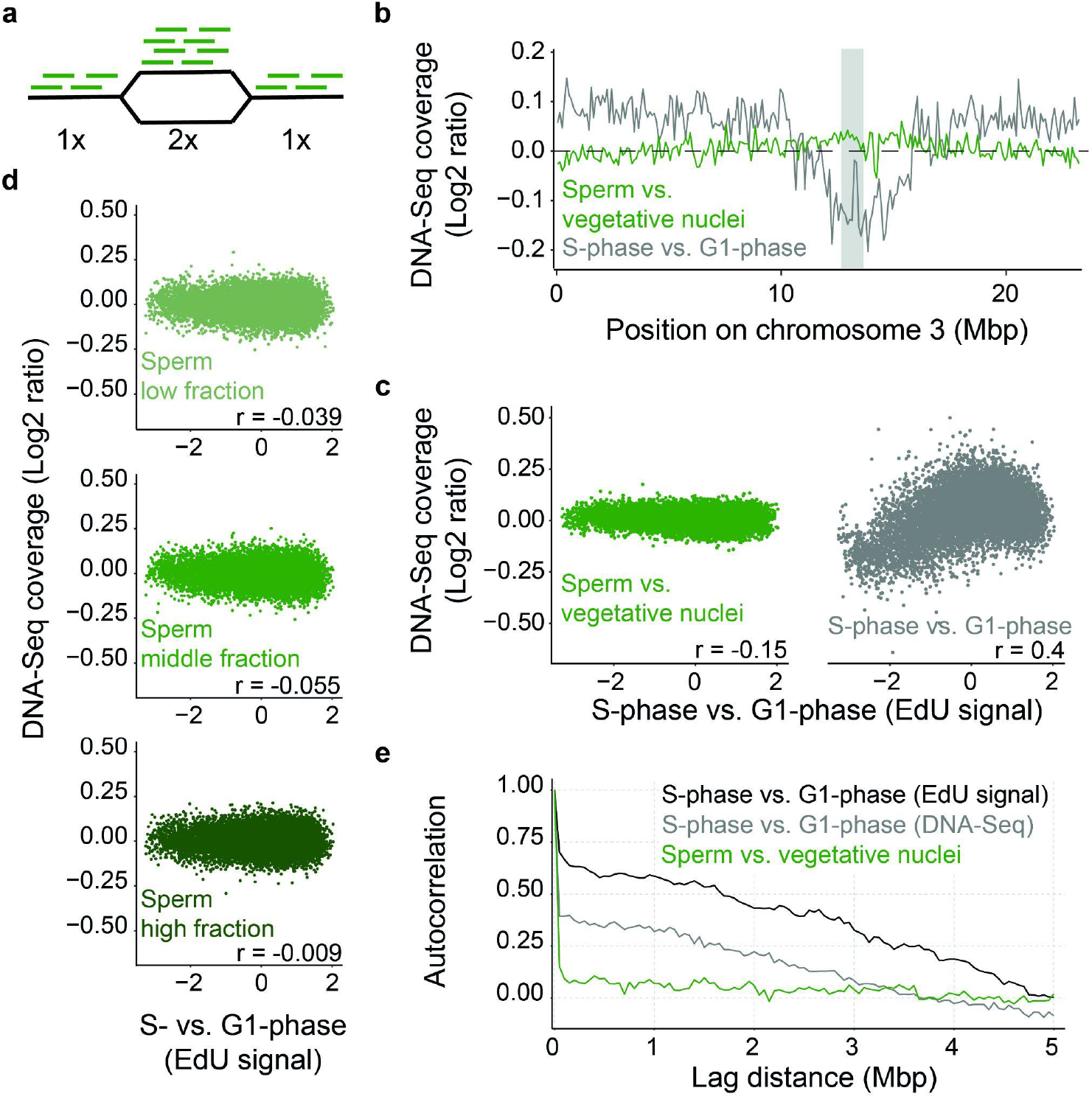
Quantification of DNA replication in sperm nuclei by DNA-Seq. **a**, Replicated regions yield double the coverage in DNA-Seq relative to non-replicated regions. **b**, DNA-Seq coverage along chromosome 3 in sperm vs. vegetative nuclei (green) or of cells enriched for S-vs. G1-phase (gray). Coverage is averaged over 100 Kbp bins and ratios are log2 transformed; shaded gray denotes the centromere position. **c**, Correlation between replication timing, as measured by EdU enrichment from early-S-phase cells^7^, to DNA-Seq coverage in sperm vs. vegetative nuclei (left) or of cells in S-phase vs. G1-phase (right). Each dot is an average of 10 Kbp bin in the genome, and Spearman correlation values are shown. **d**, Correlation of DNA-Seq coverage to replication timing as in **(c)**, for the three fractions of sperm nuclei sorted based on PI staining intensity, from lowest (top) to highest (bottom). **e**, Autocorrelation of the genomic signal to itself for different distances, for early S-vs. G1-phase enriched with EdU^7^ (black), S-vs. G1-phase quantified with DNA-Seq (gray) and sperm vs. vegetative nuclei quantified with DNA-Seq (green). The genome was averaged over 10 Kbp bins.

**Fig. 2.**
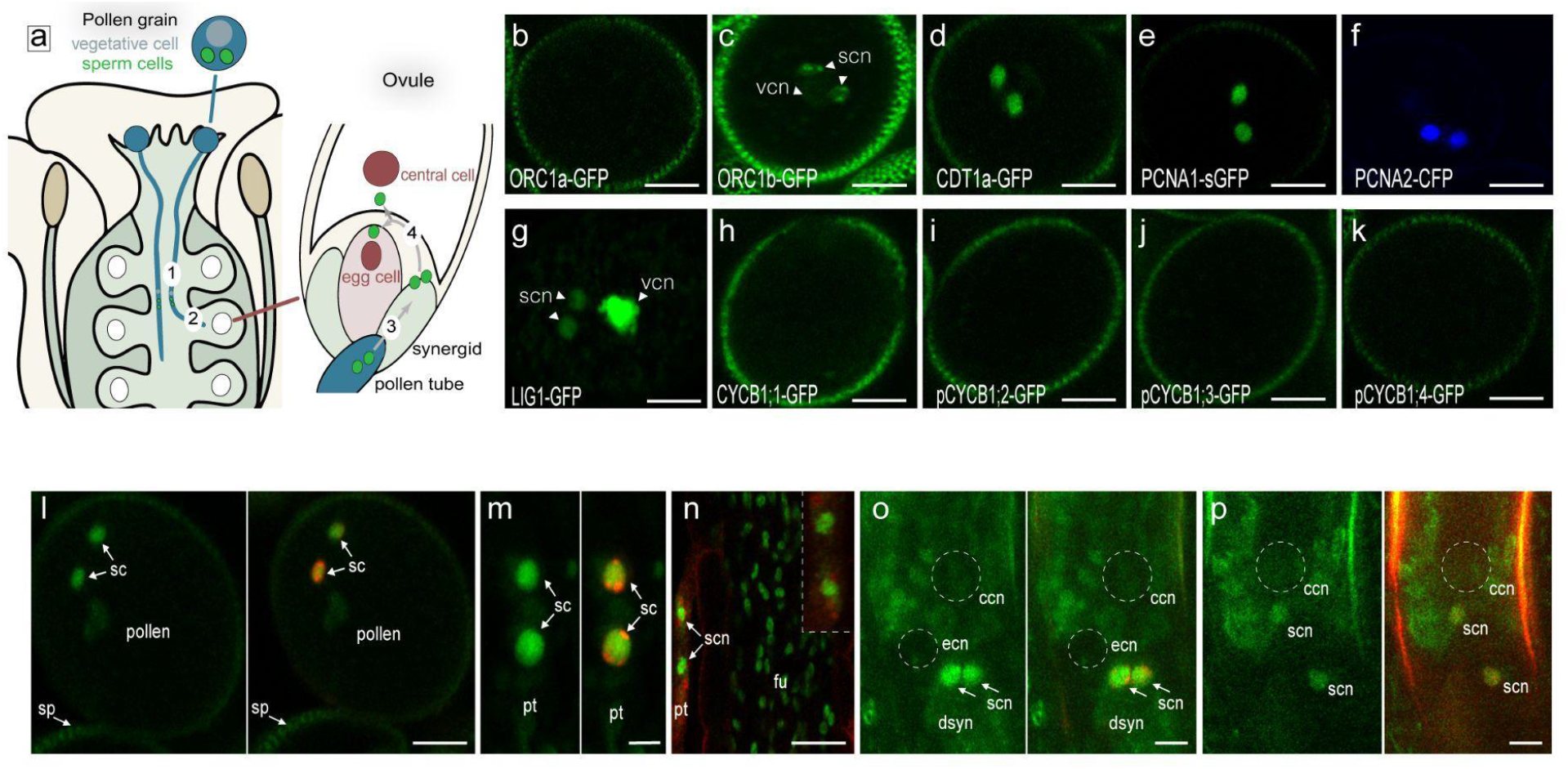
Mature sperm cells are arrested in a pre-replication phase until fertilization. **a**, Schematic representation of key steps leading to the double fertilization. Migration of sperm cells (green) in the growing pollen tube (blue) through the female sporophyte (**step 1**), along the funiculus towards the ovules (**step 2**), release of the sperm cells into the embryo sac (**step 3**), and migration towards the female gametes (egg cell and central cell, red, **step 4**). **b-k**, Expression pattern of fluorescent cell cycle phase markers in mature pollen grains. No expression of the G1 phase marker ORC1a-GFP **(b)**. Punctate fluorescent foci in sperm nuclei and weak uniform fluorescence in the vegetative cell expressing the G1 phase marker ORC1b-GFP **(c)**. The pre-replication marker CDT1a-GFP is expressed in the sperm cells **(d)**. The PCNA1-sGFP **(e)**, PCNA2-CFP **(f)**, and LIG1-GFP **(g)** markers form nuclear foci during DNA replication and a whole nuclear pattern during the G1 and G2 phases^11^. A diffuse fluorescence accumulates for the three markers in the sperm nuclei. No fluorescence is detected for all the four G2/M markers (CYCB1;1-GFP **(h)**; pCYCB1;2-GFP **(i)**; pCYCB1;3-GFP **(j)**; pCYCB1;4-GFP **(k)**. Further details on the dynamics of these cell cycle markers are provided in Extended Data Fig. 4. Number of observations n>25 for each marker. **l-p**, Dynamics of PCNA1-sGFP (green) subnuclear localization alone **(n)** or in combination with an histone 1 heterochromatin fluorescent marker (H1-1-RFP, red) **(l**,**m**,**o**,**p)** during all the key steps of sperm cell migration described in **(a). l**, Diffuse PCNA1-GFP fluorescence in sperm cell nuclei upon pollination, during pollen tube growth through the style **(m, step 1)**, along the funiculus **(n, step 2)** and after discharge in one synergid of the embryo sac **(o, step 3)**. A weaker uniform fluorescence is detected upon fertilization **(p, step 4)**. The signals around the central cell nucleus in panels **(o)** and **(p)** correspond to autofluorescence. Numbers of observations n=20 (l), n=12 (m), n= 2 (n), n= 5 (o and p). ccn, central cell nucleus; dsyn, degenerated synergid; ecn, egg cell nucleus; fu, funiculus; pt, pollen tube; sc, sperm cell; scn, sperm cell nucleus; sp, stigmatic papillae; vcn, vegetative cell nucleus. Scale bars, 10 µm **(b-k)**, 5 µm **(l-p)**.

However, we observed no correlation with published profiles of DNA replication timing, as opposed to in the control experiment (Fig. 1c). We also assessed this possibility by sorting three subpopulations of sperm cells based on increasing propidium iodide (PI) staining, which might reflect increased DNA replication (Extended Data Fig. 3a-c). Regardless of the PI fluorescence, sperm nuclei never showed a signal of DNA replication in our DNA-Seq measurements (Fig. 1d). Lastly, we also ruled out the possibility that profiles of DNA replication timing in sperm cells might differ from that published for other cell types. Genome replication profiles are assumed to be continuous, since close-by regions should replicate at similar times. Therefore, autocorrelation of the coverage between near-by positions should correlate if DNA is replicating, irrespective of the replication order. In contrast to the synchronized cells, no autocorrelation was observed for sperm cells (Fig. 1e, Extended Data Fig. 3d).

**Fig. 3.**
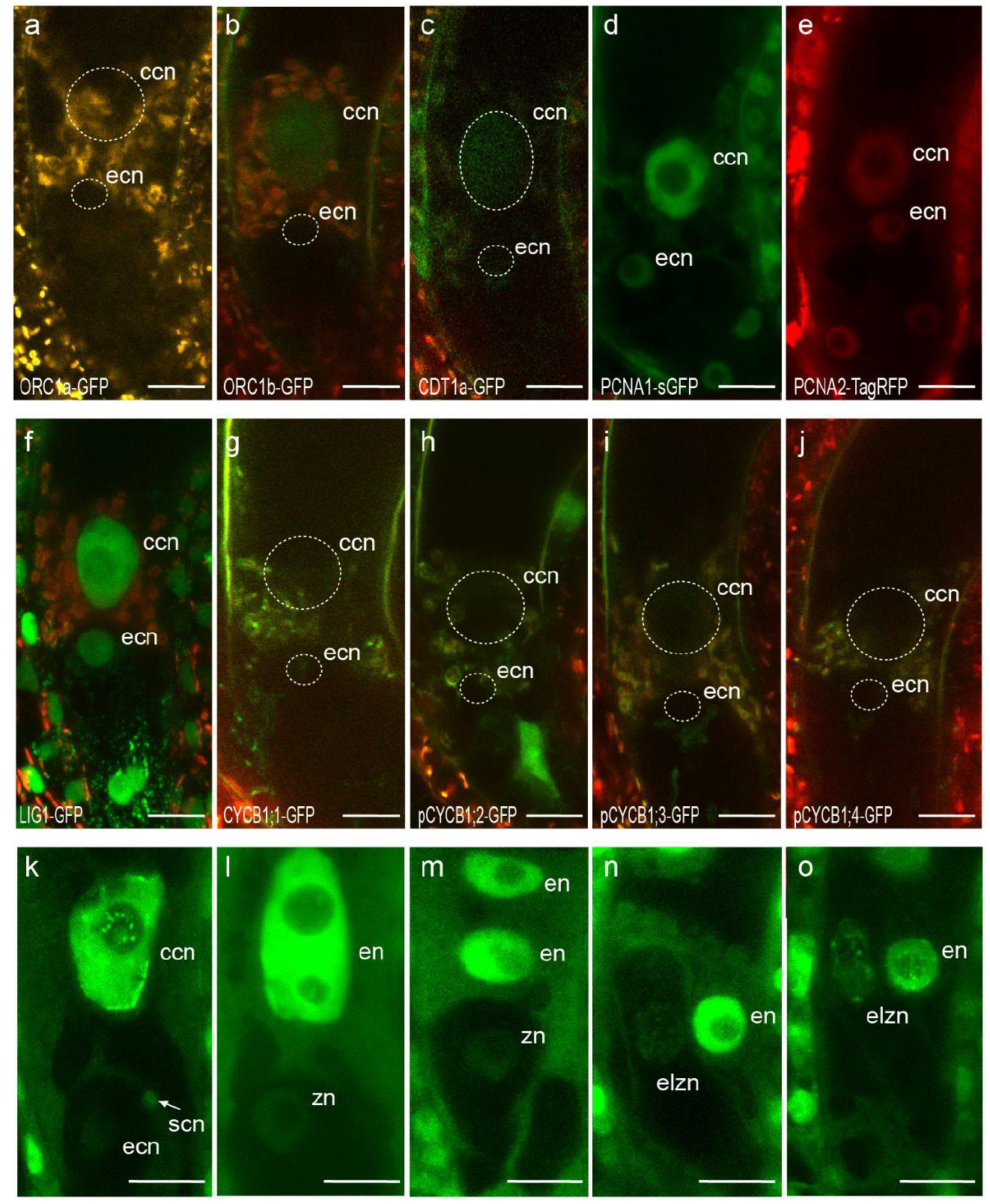
The mature egg cell and central cell are arrested in a different phase of the cell cycle before DNA replication and only start replicating at fertilization. **a-j**, Expression pattern of cell cycle phase markers in the mature female gametes. The G1-phase marker ORC1a-GFP **(a)** is not expressed in the female gametes. A weak fluorescence of the G1 phase marker ORC1b-GFP **(b)** and the pre-replication marker CDT1a-GFP is detected specifically in the central cell nucleus **(c)**. In contrast to fluorescent nuclear foci formed during the S phase^11^ (Extended Data Fig. 1), a diffuse fluorescence of the three S-phase markers PCNA1-sGFP **(d)**, PCNA2-TagRFP **(e)**, and LIG1-GFP **(f)** markers is detected in both female gametes. No expression is detected for all the four G2/M markers CYCB1;1-GFP **(g)**; pCYCB1;2-GFP **(h)**; pCYCB1;3-GFP **(i)**; pCYCB1;4-GFP **(j)**. The signals around the central-cell nucleus correspond to autofluorescence. Further details on the dynamics of these cell-cycle markers can be found in Extended Data Fig. 1. Number of observations n>25 for each marker. **k-o**, Dynamics of the PCNA1-sGFP subnuclear pattern in the developing zygote and endosperm. The egg cell and the central are fertilized by any of the two sperm cells to generate a zygote and an endosperm, respectively. **k**, The unfertilized egg cell nucleus retains a homogeneous nuclear fluorescence. The arrow indicates one sperm cell reaching the egg cell. Speckled foci of fluorescence are detected in the central cell nucleus. **l**, The zygote and the forming endosperm with two nucleoli, both show a uniform nuclear fluorescence. **m**, The elongating zygote retains a uniform nuclear fluorescence through the 2-nucleate endosperm. **n**,**o**, A speckled nuclear pattern becomes detectable in the elongated zygote through the 4-nucleate and 8-nucleate endosperm. A complete view of panels n and o is shown in Extended Data Fig. 8. Number of observations n=5 (k), n=6 (l), n=2 (m), n= 5 (n-o). ccn, central cell nucleus; ecn, egg cell nucleus; elzn, elongated zygote nucleus; en, endosperm nucleus; zn, zygote nucleus. Scale bars, 15 µm **(a-j)** and 10 µm **(k-o)**.

Thus, our genome-wide analysis could not detect any DNA replication in sperm, contrary to published results^1–3^. Our analyses suggest that DAPI-based measurement of DNA content is not a reliable method to assess cell cycle phase in sperm nuclei, probably due to their peculiar chromatin structure^4^. However, direct measurements using DNA sequencing could not determine directly if sperm cells in mature pollen were arrested in G1 or G2 phase.

Given this, we sought to determine when sperm cells do commence DNA replication. We selected Arabidopsis fluorescent marker lines expressing key components of the G1, S and G2/M phases fused with a gene encoding a fluorescent protein. Confocal analyses in root cells combining markers of G1 and S phases, of S and G2/M phases, and of the different S-phase markers revealed dynamics and subnuclear patterns that can trace distinct phases of the cell cycle (Extended Data Fig. 4). We therefore analyzed their pattern in sperm cells in mature pollen grains (Fig. 2a-k). We did not detect fluorescence for ORC1a-GFP, a component of the origin recognition complex (ORC), (Fig. 2b) in sperm cell nuclei whereas ORC1b-GFP (Fig. 2c), which is degraded at the G1/S transition in proliferating cells, showed a punctate pattern consistent with that of meristematic and early differentiating root tip cells (Vergara et al., 2023, unpublished). Likewise, ORC2-GFP, which is also acting in G1, was detected in sperm nuclei (Extended Data Fig. 5a). CDT1a, another component of the pre-replication complex (pre-RC), that accumulates in G1 and is degraded at the G1/S transition^8^, was detected in sperm cells (Fig. 2d). Proliferating cell nuclear antigen (PCNA) and DNA Ligase 1 (LIG1) are essential DNA-replication factors^9,10^. Both factors form dotted foci during early replication and speckled foci during late replication but distributed uniformly in the nucleus in G1 and G2 phases in Arabidopsis^11^ (Extended Data Fig. 4). This triphasic pattern represents highly resolute visual information for the progression of S-phase. Only a uniform fluorescence signal could be observed in sperm cells from plants expressing either of the two tagged *PCNA* genes present in the Arabidopsis genome (Fig. 2e,f) or LIG1-GFP (Fig. 2g)^12^, suggesting a cell cycle arrest in the G1 phase prior to DNA replication. To expand on the above observations, we analyzed the dynamics of the four B1-type cyclins (CYCB1) present in the Arabidopsis genome, which mark the G2/M phase^13^. As reported previously, CYC-B1;1 and CYC-B1;2 reporter lines showed no expression in sperm cells^14,15^ (Fig. 2h,i). Reporter lines of CYCB1;3 and CYCB1;4 which both include the upstream region of the gene up to the coding sequence encoding the G2/M destruction box fused to the *GFP* gene^13,16^ also showed no detectable fluorescence (Fig. 2j,k). These analyses suggest that sperm chromatin is not replicated at anthesis but is arrested in late G1 phase, in agreement with our DNA-Seq analysis. Interestingly, none of the markers tested were detected in the vegetative cell with exception of LIG1-GFP (Fig. 2c,g), with a strongly uniform nuclear fluorescence, and a faint signal of ORC1b-GFP. Our data is consistent with the vegetative cell having exited the cell cycle and being in a quiescent state^17^.

Next, we searched for PCNA1-sGFP replicating foci in sperm cell nuclei from the time of pollination until their fusion with the female gametes at karyogamy (Fig. 2a). To facilitate these observations, we crossed wild type pistils with pollen from plants expressing PCNA1-sGFP and fluorescent histone 1 (H1-1-RFP) heterochromatin markers^18^ (except for Fig. 2n where only PCNA1-sGFP marker was used). Sharp heterochromatin foci of H1-1-RFP were detectable during all phases of sperm cell migration (Fig. 2l,m,o,p). In contrast, only a diffuse fluorescent pattern of PCNA1-sGFP was observed in sperm cell nuclei during pollination and pollen tube growth (Fig. 2l-m). Semi *in vivo* pollination of wild type pistils with pollen grains expressing PCNA1-sGFP further confirmed that sperm DNA does not replicate during pollen tube growth (Extended Data Fig. 6). PCNA1-sGFP foci were still not observed upon release and migration of the sperm cells within the embryo sac (Fig. 2o,p). Although PCNA1-sGFP fluorescence becomes difficult to detect at plasmogamy (the plasma membrane fusion of gametes), we did not distinguish any foci in the sperm cells under our imaging conditions indicating that sperm cells are delivered in the G1 phase. This conclusion likely extends to many other angiosperm species that produce two sperms only during pollen tube growth leaving no sufficient time for occurrence of DNA replication.

In contrast to the sperm cells, the phase of cell-cycle arrest of female gametes (egg cell and central cell) has largely remained unexplored. Nuclear DNA content measurements suggested that both female gametes are arrested before DNA replication^19–22^. Gene expression analyses of cell cycle related genes in transcriptomes of purified Arabidopsis female gametes^23,24^ did not reveal any enrichment of cell-cycle phase-specific genes (Table S2). Female gametes are limited in number and form deep within the pistil, and therefore isolating enough for DNA-Seq is challenging. We therefore analyzed the expression pattern of our set of cell-cycle markers in the female gametes (Figs. 1a and 3). Of all the G1 markers, only weak fluorescence could be detected for ORC1b-GFP and CDT1a-GFP in the central cell (Fig. 3a-c; Extended Data Fig. 5). All three S-phase markers (LIG1-GFP, PCNA1-sGFP and PCNA2-TagRFP) accumulated nuclear fluorescence in both gametes without any detectable foci (Fig. 3d-f). G2/M markers were not detected in the female gametes (Fig. 3g-j)^25–27^. To complement these observations, we analyzed the pattern of two cell cycle fluorescent sensors in the mature female gametes^28^. The PlaCCI (Plant Cell Cycle Indicator)^29^ combines three genetic fusions - CDT1a-eCFP, CYCB1;1-YFP and H3.1-mCherry, each of which indicate the G1, G2/M and G1-to-G2/M phases, respectively. Similarly to our previous observations (Fig. 3), we only observed a weak signal of CDT1a-eCFP in the central cell using the PlaCCI markers (Extended Data Fig. 7). The second fluorescent marker reports cells in S-G2-M^30^ and contains a VENUS fluorescent protein fused to a destruction box (DB) sequence from Arabidopsis *CYCB1;1* placed under the control of a histone H4 promoter. No fluorescence was detected in the mature embryo sac (Extended Data Fig. 7). We conclude that the central cell is arrested in a pre-replication phase. Because none of the markers we used were informative in the egg cell, we propose that it is in quiescent state (G0 phase). In contrast to other eukaryotes^31,32^, no specific G0 phase markers have been described so far in plants.

To track the release of the cell-cycle block in female gametes, we performed a time-course fertilization analysis of self-pollinated plants expressing PCNA1-sGFP using the precise developmental stages previously described^33^. During plasmogamy which takes place about 1 hour after release of the gametes in the embryo sac^33^, speckled foci of PCNA1-sGFP were already evident in the elongated central cell nucleus but not in the egg cell nucleus (Fig. 3k). This PCNA1 pattern indicates late S-phase that lasts approximately 1 to 1.5 hours in somatic cells^11^. The dotted pattern corresponding to early S-phase^11^ could not be detected during our experiments. Overall, our observations suggest that DNA replication in the central cell occurs upon sperm entry as previously reported^34^. At a later stage, the fertilized central cell nucleus contained 2 nucleoli (stage F2)^33^ and showed a diffuse PCNA1-sGFP pattern. This observation is consistent with the detection of the G2/M marker CYCB1;2-YFP only 1 hour after fertilization in the endosperm nucleus^35^. A uniform to speckled pattern in the elongated zygote became apparent only once the endosperm reached the 4-to-8 nucleate stage (Fig. 3m,n,o, Extended Data Fig. 8). EdU staining experiments also indicated that replicated DNA was only detectable in zygotes accompanied by a 4 or 8-nucleate endosperm^4^.

We conclude that the cell cycle arrests before DNA replication in both male and female gametes. DNA replication only begins at fertilization, but it does so asynchronously between the zygote and the endosperm. Importantly the zygote undergoes reprogramming of parental epigenetic patterns involving histone H3 gene family^36^ but also a maternal-to-zygotic transition leading to a rapid degradation of maternal transcripts and large-scale *de novo* zygotic transcription^23^. Design of gamete-specific and brighter DNA replication reporter lines will help to further refine whether these genetic and epigenetic events take place concomitantly with DNA replication during fertilization.

## Acknowledgements

We thank the following labs for sharing marker lines; A. Schnittger (CYCB1 gene reporters), U. Grossniklaus (ORC2-GFP), C. Baroux (H1-1-RFP), S. Matsunaga (PCNA1-sGFP), J. Murray (pH4-DB-CYCB1;1-Venus). We are grateful to P. Bourguet for connecting the dots. M.I. and C.M. were supported by the French National Research Agency (ANR-15-CE12-0012; ANR-21-CE20-0047). Y.V was supported by a Marie Skłodowska-Curie individual fellowship (101028014). C.G was supported by RTI2018-094793-B-I00 and PID2021-123319NB-I00 by the Ministerio de Ciencia e Innovación (MICIN and FEDER), and by ERC-AdG2018-833617 by the European Union. This project has received funding from Austrian Science Fund (FWF) including the Lise Meitner program to M.B. M1818 and grant I2363 B16 to F.B and core funding from the Austrian Academy of Sciences to F.B. and A.S. We would like to thank the Plant Sciences and Next Generation Sequencing at the Vienna BioCenter Core Facilities (VBCF), and the Biooptics facilities at IMBA/IMP. We acknowledge the imaging facility MRI, member of the National Infrastructure France-BioImaging Infrastructure supported by the French National Research Agency (ANR-10-INBS-04, Investments for the future).

## Author contributions

M.I and Y.V designed the study and wrote the manuscript. M.I and C.M. performed the microscopic experiments. B.H, Y.V, V.N performed FACS purification and DNA-Seq experiments. A.S performed the MM2d synchronization experiment. Z.V, B.D and C.G provided pre-RC-GFP lines and input on pre-RC dynamics during the cell cycle. B.J, M.B, F.B and M.N provided valuable input in the conceptualization of the study.

## Data availability

All sequencing data was deposited to SRA under accession number PRJNA914255.

**Extended Data Fig. 1.**
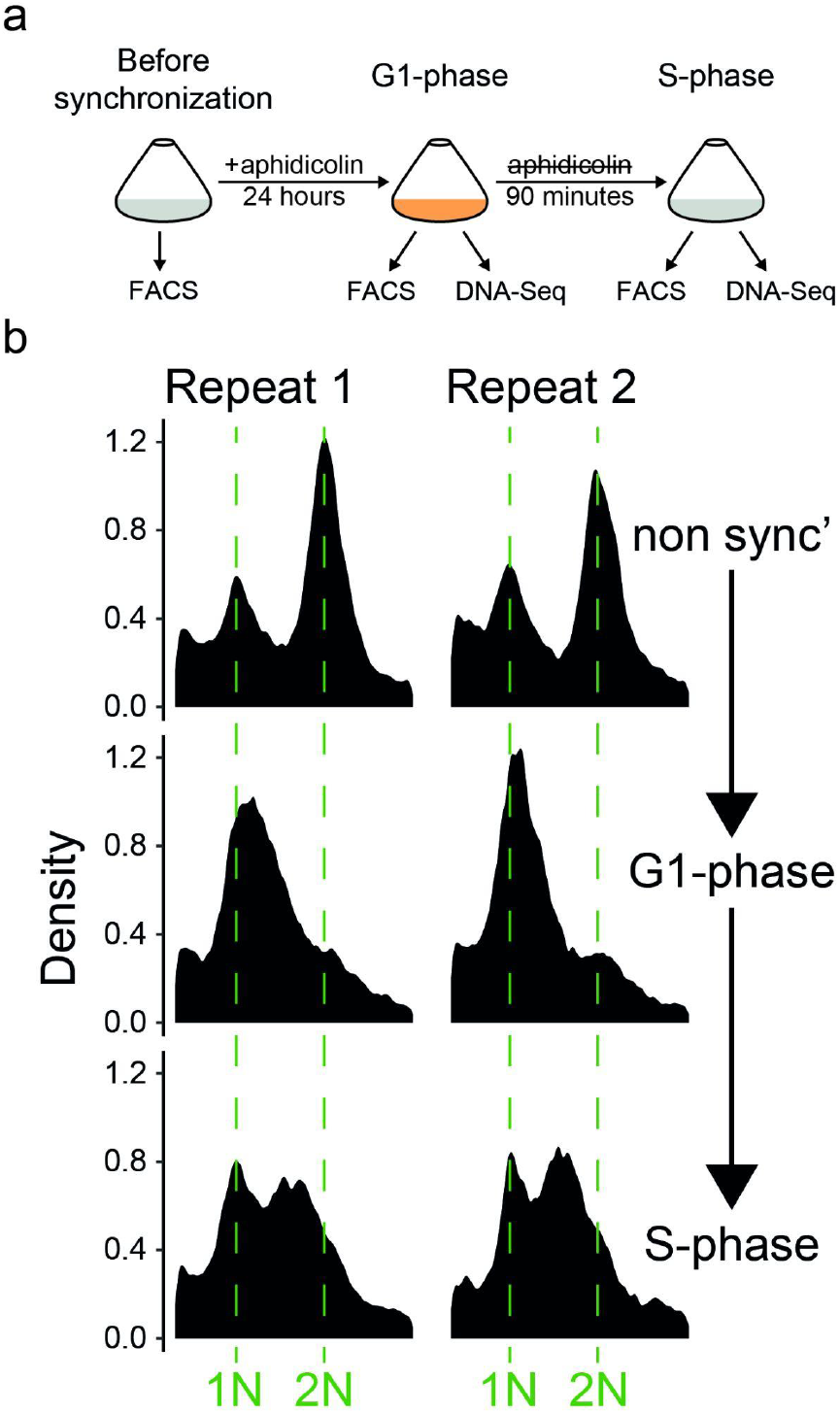
Synchronizing of cell suspension to G1 and release into S-phase. **a**, *A. thaliana* suspension cells were grown at room temperature in the dark, with shaking. These cells were synchronized to the G1 phase of the cell cycle by incubating them with aphidicolin for 24 hours. After this period, the aphidicolin was removed, allowing the cells to re-enter the cell cycle. Samples were collected at three different time points: before synchronization, after synchronization, and 90 minutes after the release from synchronization. At each time point, samples were collected for flow cytometry analysis, and DNA sequencing was performed on the samples collected at the G1- and S-phase time points. **b**, FACS profiles of DNA content measurements as quantified by DAPI staining shown at the three time points described in (**A**) for the two replicates of the experiment.

**Extended Data Fig. 2.**
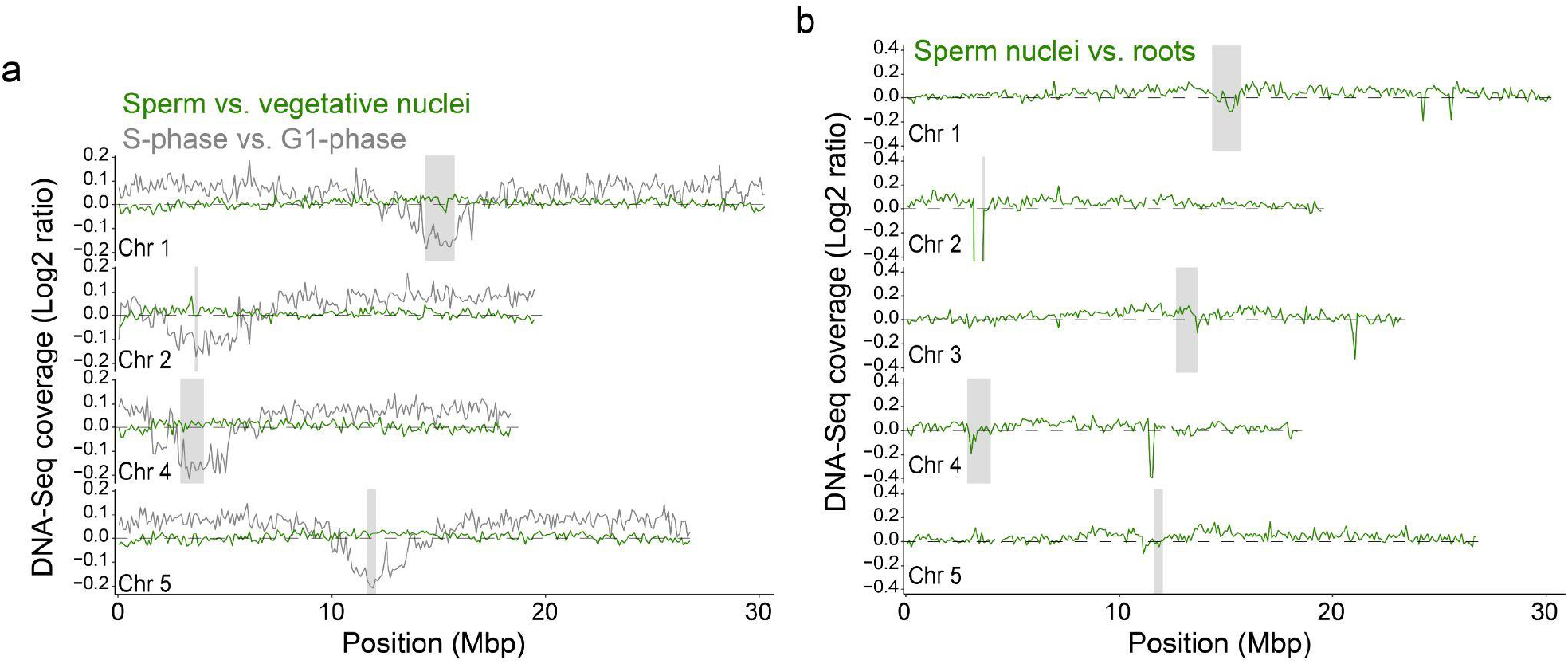
DNA-Seq coverage along *A. thaliana* chromosomes. **a**, DNA-Seq coverage as in Fig. 1b along chromosomes 1,2,4,5. **b**, The DNA-Seq coverage of sperm nuclei, normalized by DNA-Seq from root nuclei <u>^37^</u>, is shown for all five chromosomes of *A. thaliana* as in **(a)**.

**Extended Data Fig. 3.**
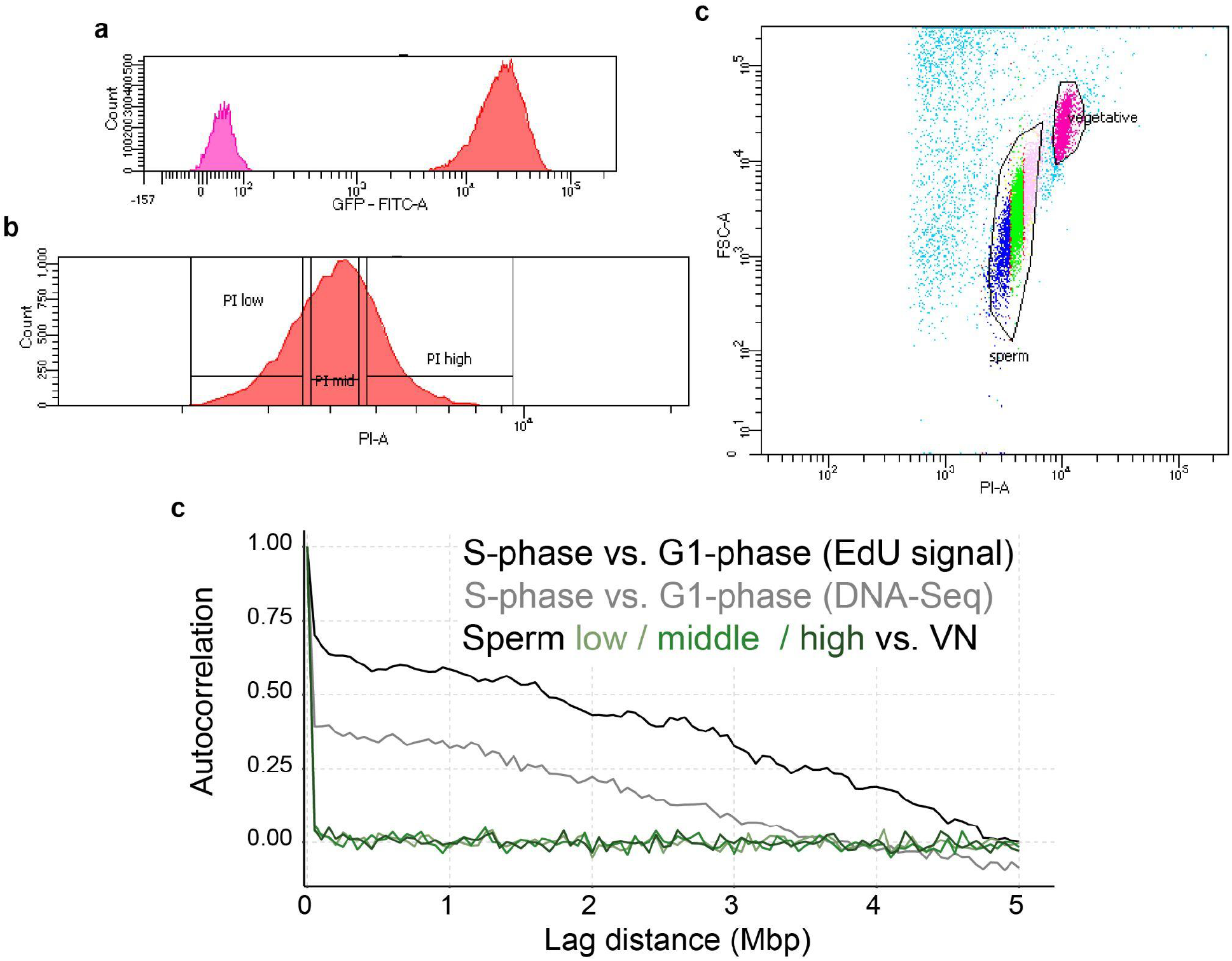
FACS sorting of sperm nuclei to three fractions. The sperm and vegetative nuclei in the HTR10-Clover sample were initially distinguished based on the Clover (GFP in axis) fluorescence (**a**). The sperm nuclei were then further divided into three fractions based on the DNA dye propidium iodide (PI), as shown in (**b**). The final sorting configuration, using PI, Clover, and forward scatter (FSC), is depicted in (**c**). **d**, The three fractions of sperm nuclei’s autocorrelation are shown in different shades of green as shown in Fig. 1e. The S-phase vs. G1-phase autocorrelations are once more displayed for comparison, as in Fig. 1e.

**Extended Data Fig. 4.**
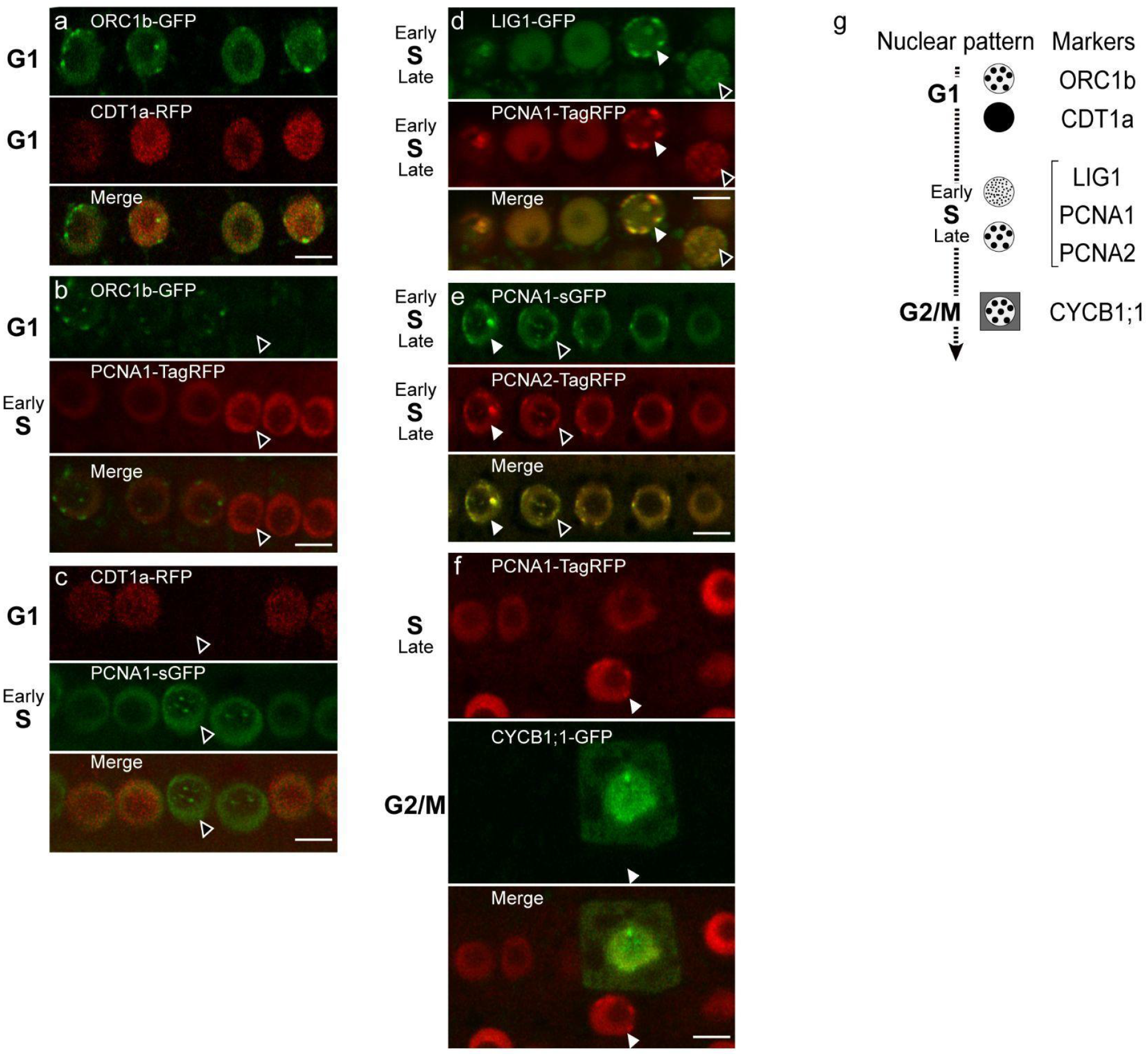
Dynamics of representative cell-cycle markers used in this study in root cells. Confocal images were obtained from cells of the root meristematic zone expressing a combination of two cell cycle phase markers. These markers all comprise the entire gene fused to a gene encoding a fluorescent protein. **a**, ORC1b-GFP; CDT1a-RFP. **b**, ORC1b-GFP; PCNA1-TagRFP. **c**, CDT1a-RFP; PCNA1-sGFP. **d**, LIG1-GFP; PCNA1-TagRFP. **e**, PCNA1-sGFP; PCNA2-TagRFP. **f**, PCNA1-TagRFP; CYCB1;1-GFP. Empty and filled arrowheads indicate S phase nuclei with dotted and speckled foci, respectively. They correspond to early and late S phase nuclei, respectively. n>15 for each combination of markers. Scale bars, 10 µm. **g**, Diagram summarizing the typical nuclear patterns of the representative cell cycle phase markers used in this study. Circles filled with dots represent a focalized nuclear pattern. For the S-phase markers, circles filled with small dots or large dots indicate a dotted or speckled nuclear pattern, respectively. Filled black circles indicate a homogeneous nuclear pattern. A gray square symbolizes cytoplasmic accumulation.

**Extended Data Fig. 5.**
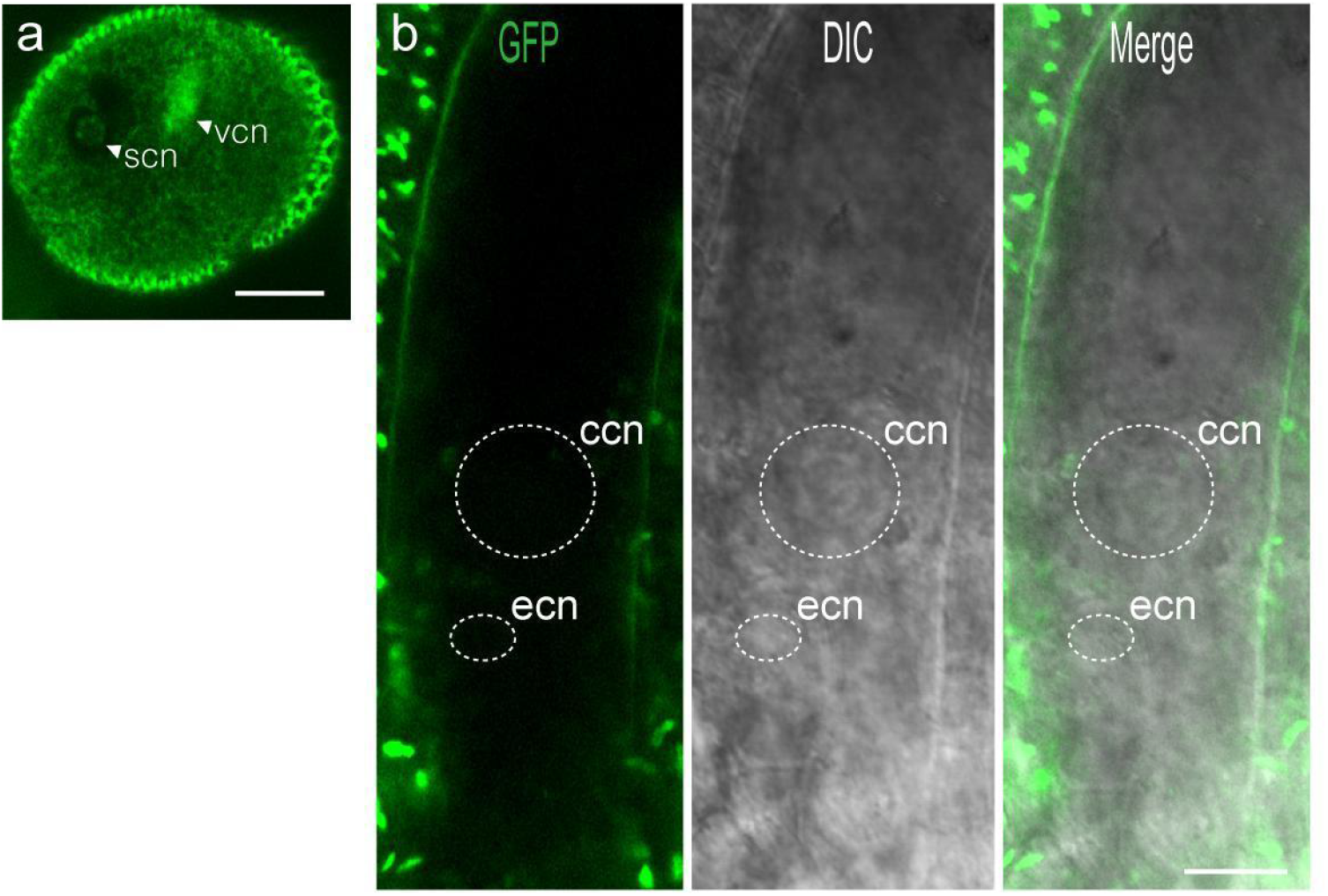
Expression pattern of ORC2-GFP translational fusion in the mature pollen grain and embryo sac. Confocal images were obtained from mature pollen grains **(a)** and embryo sacs **(b)** of plants expressing pORC2-ORC2-GFP<u>^38^</u>. **a**, A punctate fluorescent signal is detected in the sperm nuclei (n=35). **b**, No fluorescence is detected in the mature embryo sac (n=25). ccn, central cell nucleus; ecn, egg cell nucleus; scn, sperm cell nucleus; vcn, vegetative cell nucleus. Scale bars 10 µm **(a)** and 15 µm **(b)**.

**Extended Data Fig. 6.**
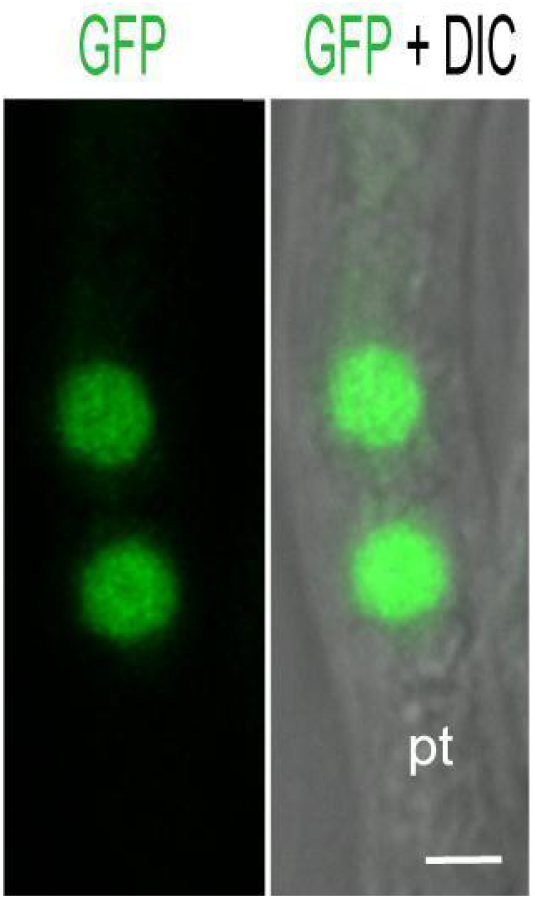
Subnuclear pattern of PCNA1-sGFP in sperm cells in growing pollen tubes. Wild type pistils were pollinated with pollen grains expressing the pPCNA1-PCNA1-sGFP reporter and cut at the end of the style. Confocal images were taken from growing pollen tubes (pt) emerging from the style for 4 h after pollination (n=45). A uniform fluorescent signal is detected in the sperm nuclei. Scale bar, 10 µm.

**Extended Data Fig. 7.**
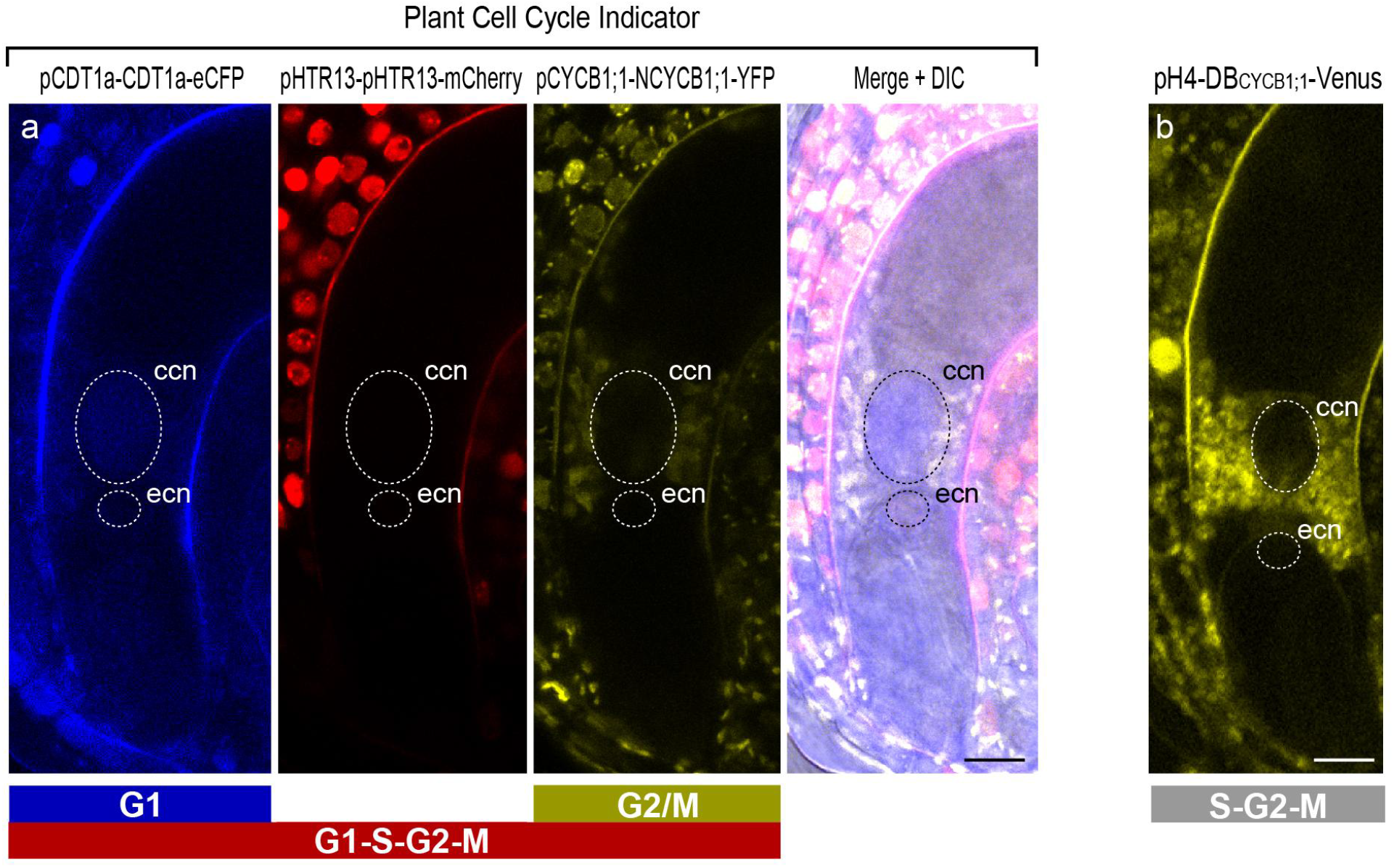
Dynamics of cell-cycle reporters in the egg cell and the central cell in the mature embryo sac. Confocal images obtained from mature embryo sacs from transgenic plants expressing either the Plant Cell Cycle Indicator sensor^29^ **(a)** or the S-G2-M phase fluorescent reporter (pH4-DB^CYCB1;1^-Venus)^30^ **(b). a**, A weak fluorescence of CDT1a-eCFP was detected in the central cell nucleus (n=15). None of the cell cycle phase markers were detected in the egg cell nucleus. **b**, No fluorescence was detected in the egg cell and central cell (n=25). The signals around the central cell nucleus correspond to autofluorescence. ccn, central cell nucleus; ecn, egg cell nucleus. Scale bars, 15 µm.

**Extended Data Fig. 8.**
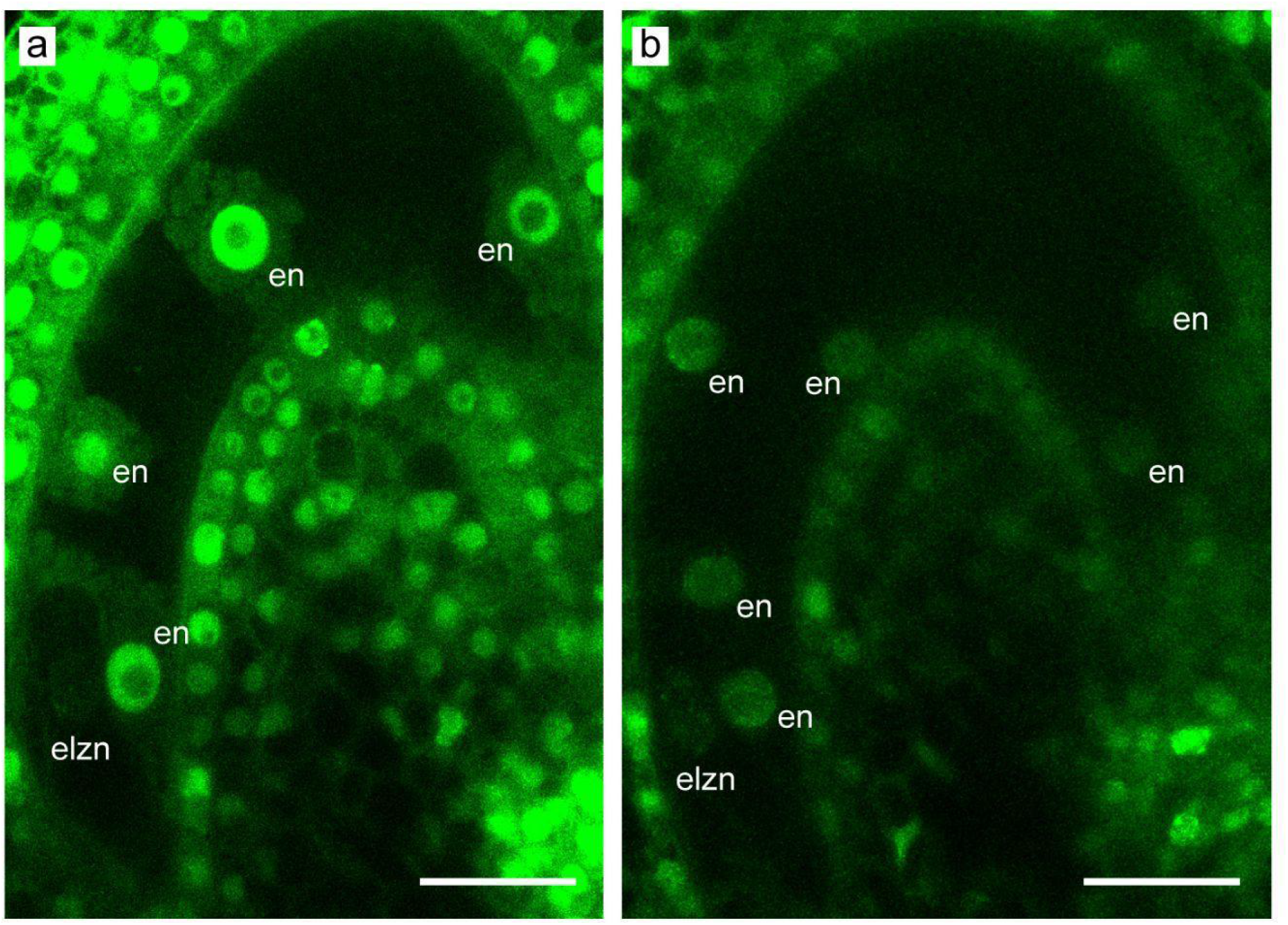
Subnuclear pattern of PCNA1-sGFP in the developing zygote. Confocal images were generated from transgenic plants expressing pPCNA1-PCNA1-sGFP. At the 4-nuclear **(a, n=5)** and 8-nuclear endosperm stage **(b, n=5)**, the zygote becomes elongated and speckled foci of PCNA1-GFP are detected in the nucleus.. elzn, elongated zygote nucleus; en, endosperm nucleus. Scale bars, 20 µm.

## Methods

### Plant materials and growth conditions

All *A. thaliana* plants described in this study were in the Col-0 background (except when mentioned). The following reporter lines were described previously: pORC2-ORC2-GFP^38^ (Ler accession), pORC1a-ORC1a-GFP and pORC1b-ORC1b-GFP (Vergara et al. unpublished), pLIGI-LIGI-GFP^12^, pHTR10-HTR10-Clover^6^, pPCNA1-PCNA1-sGFP^11^, pH4-DB-CYCB1;1-Venus^30^, pCDT1a-CDT1a-GFP^8^, pH1-1-H1-1-RFP^18^, Plant CCI^29^. Reporter lines for all four *CYCB1* genes were provided by Arp Schnittger^13^. After a 4 day-stratification in the dark, seeds were germinated and grown on soil in a growth chamber under long days at 20°C (16-h light/8-h night). For the purpose of collecting pollen for FACS, seeds were sown in 60 cm x 40 cm trays with 6 cm of soil (4:1 Gramoflor 2006:perlite), stratified for 4 days at 4°C in the dark, and then moved to 21°C/16°C with 60% humidity, 16-h light/8-h night.

### Cloning and plant transformation

A PCNA2 genomic fragment comprising 412 bp upstream of the ATG until the last codon before the termination codon of the gene flanked with attL Gateway recombination sites was generated by gene synthesis (Genscript). The pPCNA1-PCNA1-TagRFP reporter construct was obtained after a LR Clonase II reaction using the pPCNA1-PCNA1 entry vector^11^ and the destination vector pGWB559^39^. Similarly the pPCNA2-PCNA2-TagRFP and pPCNA2-PCNA2-CFP constructs were generated using pGWB459 and pGWB543 destination vectors^39^, respectively. The transgenic plants were generated by floral dipping^40^ and selected on MS solid medium (Duchefa) with the appropriate selective agent.

### Sample preparation and microscopy

Observations of the different reporter lines during male and female gametogenesis were obtained from freshly dissected anthers and carpels. Self-pollinated pistils expressing the PCNA1-sGFP marker^11^ or pistils pollinated with the PCNA1-sGFP and H1-1-RFP^18^ markers were dissected and mounted on ½ MS supplemented with 0.4% Phytagel. All the steps ranging from pollination, growth of the pollen tubes in the pistil up to the double fertilization process and consequent development of the fertilization products (zygote, endosperm) were reconstituted from a series of pistils dissected at distinct time after pollination ^36^. Semi–i*n vivo* pollen tube growth was performed as described before^41^. Wild type pistils were pollinated with pollen grains expressing the PCNA1-sGFP marker and cut at the end of the style. The truncated pistils were incubated on a pollen tube germination medium at 25°С for 4 h. Pollen tube growth was observed 4 to 6 h after pollination. Imaging of cell cycle reporters in roots was performed as previously described^42^.

All the images were obtained using a laser scanning confocal microscope (Zeiss LSM880 Fast Airyscan). Image contrast and brightness were adjusted with Adobe Photoshop and processed images were assembled in Adobe Illustrator.

### Pollen collection and Fluorescence-activated cell sorting (FACS)

Pollen was harvested from open flowers using the vacuum suction method^43^, using 150 μm, 60 μm and 10 μm mesh filters and then flash freezed and kept in -80°C. Sperm and vegetative nuclei were isolated by FACS following the previously described method^44^. Collected pollen was hydrated by resuspension in an ice-cold Galbraith buffer (45 mM MgCl_2_, 30 mM Tri-Sodium Citrate, 20 mM MOPS pH 7.0, 0.1% Triton X-100) with added 72 mM ß-mercaptoethanol and 1× complete protease inhibitor cocktail (Roche)^45^. Pollen suspension was then centrifuged for 1 min at 10,000 g, 50 μl of 100 μm glass beads were added and vortexed for 4 min to disrupt pollen walls. The suspension was filtered through a 10 μm nylon mesh to retrieve the sperm and the vegetative nuclei. Nuclei were then stained with SYBR Green (S9430, Sigma-Aldrich) or propidium iodide (P4170, Sigma-Aldreich).

A BD FACS AriaTM III Cell Sorter was used to sort the nuclei suspension. The sorter was operated according to standard configuration, utilizing a 70 μm ceramic nozzle with a 1 x PBS solution flowing at a constant 20 psi pressure. Sperm and vegetative nuclei were collected in two different tubes. For two replicates of the experiment, HTR10-Clover sperm nuclei were further separated in the FACS to three groups based on their propidium iodide fluorescence. The different experiments and sequenced samples are summarized in table S1.

### S-phase synchronization and release of MM2d cell suspension culture

MM2d cells from *Arabidopsis thaliana* (Ler) were obtained from the lab of Crisanto Guiterrez and were previously described^46,47^. Cells were grown at room temperature in the dark shaking at 130 rpm and subcultured every week by adding 3 ml of the old culture into 50 ml fresh MSS media (1× MS, 3% sucrose 0.5 mg/l NAA, 0.05 mg/l kinetin). For cell cycle synchronization, one-week-old MM2d cells were diluted in new MSS media in a 1:5 ratio. After taking a sample for flow cytometry analysis, 4 μg/ml of Aphidicolin (Sigma) was added to the diluted culture for the cell-cycle block. Cells were incubated for 24 h with shaking at 130 rpm at room temperature in the dark. After 24 h, a sample was taken for flow cytometry analysis and 40 ml of cells were harvested in liquid nitrogen. Cells were washed with MMS media to remove the Aphidicolin and were resuspended in media to resume the cell cycle synchronously. After 90 min, when cells are enriched for S-phase, a sample was taken for flow-cytometry analysis, and another 40 ml were harvested in liquid nitrogen. The experiment was done in two biological replicates. For flow cytometry analysis, samples were chopped in Galbraith buffer (45 mM MgCl2, 30 mM Nacitrate, 20 mM MOPS pH 7.0, 0.1% Triton X-100) and filtered through a 40 μm nylon mesh. Nuclei were stained with 4 ug/ml DAPI and analyzed on the Penteon flow cytometer. FACS profiles were analyzed using flowCore^48^. To harvest the samples of G1 and 90 min following release into S phase, cells were filtered using a cell strainer, dried with filter sheets, and then flash frozen in liquid nitrogen. To break the cells, Precellys Zirconium oxide beads were added to the frozen cell pellets. Cells were disrupted by shaking 3 times at 5800 rpm for 20 s using the Precellys Evolution machine. Cells were frozen in liquid nitrogen between each disruption. DNA extraction, sonication of DNA and construction of libraries for next-generation sequencing was done as for the sperm and vegetative nuclei.

### Isolation of DNA from nuclei and library preparation

Genomic DNA was extracted using QIAamp DNA Micro Kit (56304) following the provider “Small Volumes of Blood’’ protocol. DNA was diluted to 10 ng in 50 μl of an elution buffer (10 mM Tris-HCL, pH 8.0) in microTube AFA Fiber tubes (520052). Samples were sonicated in Covaris E220 programmed to 80 s / 10 Duty / 140 PIP / 200 cycles. 25 μl of the sonicated DNA was used for library preparation using the NEBNext® Ultra™ II DNA Library Prep Kit for Illumina (E7645S), with a final amplification of 12 PCR cycles. Libraries were sequenced on Illumina Novaseq or Nextseq machines, with at least 10 million paired-end reads per library.

### Processing of DNA-Seq libraries

All next-generation sequencing data used in this work were processed in exactly the same way, including the previously published datasets. To avoid detecting effects attributed to technical differences in sequencing mode and coverage, sequence reads were down-sampled to 10^7^ paired-reads using seqtk v1.3, and trimmed to 50 bp for both R1 and R2 using Trim Galore v0.6.2 (--hardtrim5 50). Reads were then further trimmed using Trim Galore v0.6.2 (--paired --fastqc). Reads were then mapped to the TAIR10 genome using bowtie2 v2.3.5.1 with default parameters^49^. Aligned reads were converted to indexed-sorted bam using samtools v1.10^50^ and duplicate reads were filtered out using Picard v2.18.27 MarkDuplicates option. Coverage per 10kb or 100kb bins of the genome were calculated using DeepTools v3.3.1 (-of bedgraph)^51^. The procedure described above was wrapped in a Nextflow pipeline^52^, and was uploaded to GitHub. Genomic coverage per sample per bin size was then normalized to the same total signal and log2 transformed. Previously published data on genomic DNA from Col-0 roots^37^ was downloaded from SRA, accession number SRR1945757, and data for early, middle, and late S-phase cells labeled with EdU along with G1-phase cells^7^ were downloaded from SRR3931891 - SRR3931900.

---

**Table S1.**
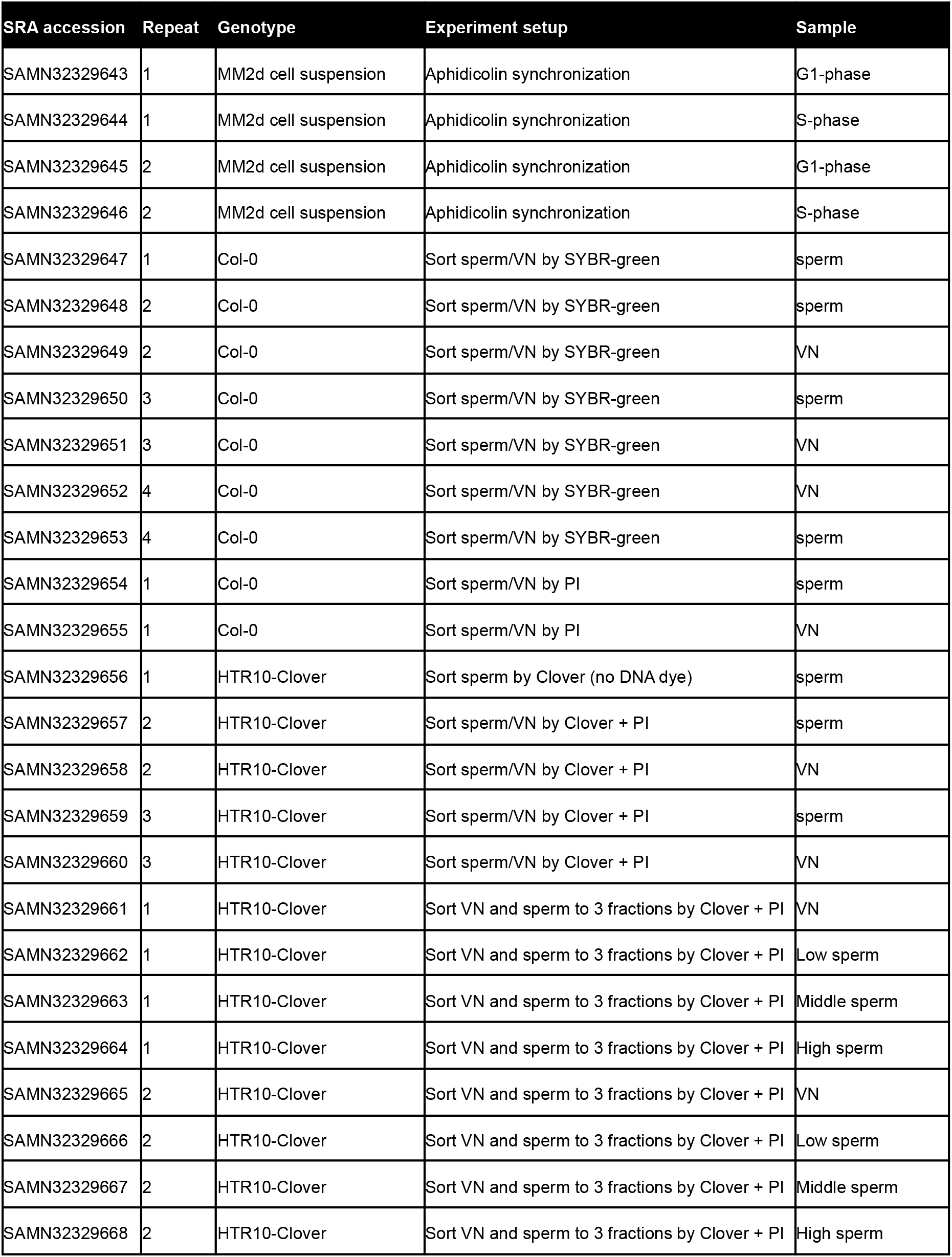
List of sequenced samples

| **SRA accession** | **Repeat** | **Genotype** | **Experiment setup** | **Sample** |
| --- | --- | --- | --- | --- |
| SAMN32329643 | 1 | MM2d cell suspension | Aphidicolin synchronization | G1-phase |
| SAMN32329644 | 1 | MM2d cell suspension | Aphidicolin synchronization | S-phase |
| SAMN32329645 | 2 | MM2d cell suspension | Aphidicolin synchronization | G1-phase |
| SAMN32329646 | 2 | MM2d cell suspension | Aphidicolin synchronization | S-phase |
| SAMN32329647 | 1 | Col-0 | Sort sperm/VN by SYBR-green | sperm |
| SAMN32329648 | 2 | Col-0 | Sort sperm/VN by SYBR-green | sperm |
| SAMN32329649 | 2 | Col-0 | Sort sperm/VN by SYBR-green | VN |
| SAMN32329650 | 3 | Col-0 | Sort sperm/VN by SYBR-green | sperm |
| SAMN32329651 | 3 | Col-0 | Sort sperm/VN by SYBR-green | VN |
| SAMN32329652 | 4 | Col-0 | Sort sperm/VN by SYBR-green | VN |
| SAMN32329653 | 4 | Col-0 | Sort sperm/VN by SYBR-green | sperm |
| SAMN32329654 | 1 | Col-0 | Sort sperm/VN by PI | sperm |
| SAMN32329655 | 1 | Col-0 | Sort sperm/VN by PI | VN |
| SAMN32329656 | 1 | HTR10-Clover | Sort sperm by Clover (no DNA dye) | sperm |
| SAMN32329657 | 2 | HTR10-Clover | Sort sperm/VN by Clover + PI | sperm |
| SAMN32329658 | 2 | HTR10-Clover | Sort sperm/VN by Clover + PI | VN |
| SAMN32329659 | 3 | HTR10-Clover | Sort sperm/VN by Clover + PI | sperm |
| SAMN32329660 | 3 | HTR10-Clover | Sort sperm/VN by Clover + PI | VN |
| SAMN32329661 | 1 | HTR10-Clover | Sort VN and sperm to 3 fractions by Clover + PI | VN |
| SAMN32329662 | 1 | HTR10-Clover | Sort VN and sperm to 3 fractions by Clover + PI | Low sperm |
| SAMN32329663 | 1 | HTR10-Clover | Sort VN and sperm to 3 fractions by Clover + PI | Middle sperm |
| SAMN32329664 | 1 | HTR10-Clover | Sort VN and sperm to 3 fractions by Clover + PI | High sperm |
| SAMN32329665 | 2 | HTR10-Clover | Sort VN and sperm to 3 fractions by Clover + PI | VN |
| SAMN32329666 | 2 | HTR10-Clover | Sort VN and sperm to 3 fractions by Clover + PI | Low sperm |
| SAMN32329667 | 2 | HTR10-Clover | Sort VN and sperm to 3 fractions by Clover + PI | Middle sperm |
| SAMN32329668 | 2 | HTR10-Clover | Sort VN and sperm to 3 fractions by Clover + PI | High sperm |

**Table S2.** Expression of selected DNA replication genes in transcriptome of purified egg cell and central cell of Arabidopsis.

|  |  | Egg cell |  |  |  | Central cell |  |
| --- | --- | --- | --- | --- | --- | --- | --- |
| Gene Description | Gene ID | EC | EC1 | EC2 | EC3 | CC1 | CC2 |
|  |  | Zhao et al. 2019* | Susaki et al. 2021** |  |  | Susaki et al. 2021** |  |
| Prereplication complex assembly |  |  |  |  |  |  |  |
| ORC1A | At4g14700 | 2.23 | 0,1 | 2,2 | 0,0 | 1,8 | 15,8 |
| ORC1B | At4g12620 | 0,0 | 5,5 | 22,4 | 5,1 | 14,4 | 16,8 |
| ORC2 | At2g37560 | 29,0 | 4,3 | 74,3 | 15,2 | 74,5 | 73,5 |
| ORC3 | At5g16690 | 3,1 | 3,4 | 0,0 | 0,0 | 19,0 | 30,3 |
| ORC4 | At2g01120 | 26,4 | 56,1 | 30,6 | 69,4 | 116,3 | 90,7 |
| ORC5 | At4g29910 | 27,1 | 1,6 | 0,0 | 38,5 | 71,3 | 36,8 |
| ORC6 | At1g26840 | 0,7 | n.d. | n.d. | n.d. | n.d. | n.d. |
| NOC3 | At1g79150 | 40,4 | 121,8 | 134,0 | 90,6 | 93,8 | 63,9 |
| Cdc6a | At2g29680 | 0,00 | 0,0 | 0,0 | 0,0 | 48,8 | 96,0 |
| Cdc6b | At1g07270 | 1,49 | 0,0 | 0,0 | 0,0 | 34,0 | 15,5 |
| Cdt1a | At2g31270 | 8,10 | 49,1 | 43,1 | 0,0 | 149,0 | 115,7 |
| Cdt1b | At3g54710 | 49.52 | 30,1 | 7,8 | 1,5 | 9,7 | 44,6 |
| Cdc45 | At3g25100 | 1,8 | 20,8 | 3,8 | 10,8 | 1,1 | 23,3 |
| Mcm9 | At2g14050 | 6,6 | 5,7 | 58,5 | 7,3 | 36,2 | 12,2 |
| Replication fork assembly |  |  |  |  |  |  |  |
| Mcm8 | At3g09660 | 3,1 | 3,5 | 0,0 | 11,5 | 22,2 | 8,5 |
| Mcm10 | At2g20980 | 0,0 | 20,4 | 6,9 | 0,0 | 79,6 | 79,0 |
| TOPBP | At1g77320 | 14,2 | 100,1 | 1,2 | 59,7 | 16,6 | 34,0 |
| Psf1 | At1g80190 | 0,8 | 0,0 | 0,0 | 0,0 | 108,9 | 30,4 |
| Psf2 | At3g12530 | 7,05 | 0,0 | 0,0 | 0,0 | 20,6 | 71,6 |
| Psf3 | At1g19080 | 27,7 | 69,0 | 94,4 | 108,0 | 144,1 | 37,8 |
|  | At3g55490 | 42,9 | 69,2 | 156,8 | 73,1 | 109,5 | 63,5 |
| SLD5 | At5g49010 | 7,0 | 5,5 | 0,4 | 4,5 | 4,0 | 3,1 |
| DNA synthesis at replication forks |  |  |  |  |  |  |  |
| POLA1 | At5g67100 | 4,2 | 6,3 | 6,6 | 1,8 | 101,8 | 166,6 |
| POLA2 | At1g67630 | 0,0 | 11,2 | 0,0 | 0,0 | 108,5 | 23,1 |
| POLA3 | At1g67320 | 7,9 | 30,4 | 16,9 | 17,0 | 99,2 | 67,0 |
| POLA4 | At5g41880 | 1,3 | 0,0 | 0,0 | 0,0 | 99,7 | 40,4 |
| POLD1 | At5g63960 | 6,2 | 41,0 | 15,5 | 4,0 | 49,2 | 82,1 |
| POLD2 | At2g42120 | 57,5 | 10,6 | 10,9 | 8,2 | 7,3 | 88,0 |
| POLD3 | At1g78650 | 26,8 | 68,1 | 127,6 | 69,7 | 95,6 | 35,3 |
| POLD4 | At1g09815 | 37,1 | 29,7 | 0,0 | 4,9 | 0,0 | 57,4 |
| POLE1 | At1g08260 | 2,5 | 19,7 | 11,9 | 5,8 | 83,1 | 35,0 |
| POL2B | At2g27120 | 2,5 | 25,1 | 9,0 | 17,5 | 59,4 | 44,4 |
| POLE2 | At5g22110 | 2,7 | 11,6 | 0,0 | 0,0 | 0,0 | 116,8 |
| RPA1a | At2g06510 | 22,47 | 85,5 | 4,6 | 11,3 | 4,1 | 26,3 |
| RPA1b | At5g08020 | 0,25 | 0,0 | 0,0 | 0,0 | 7,4 | 2,5 |
| RPA1c | At5g45400 | 50,4 | 62,2 | 59,1 | 97,4 | 30,2 | 75,5 |
| RPA1d | At5g61000 | 66,1 | 91,6 | 81,1 | 57,8 | 89,8 | 40,3 |
| RPA1e | At4g19130 | 13,1 | 39,6 | 22,3 | 10,9 | 13,8 | 38,6 |
| RPA2a | At2g24490 | 17,8 | 36,3 | 0,0 | 28,2 | 43,3 | 45,5 |
| RPA2b | At3g02920 | 108,6 | 421,0 | 1101,8 | 253,9 | 79,3 | 149,8 |
| RPA3a | At3g52630 | 0,0 | 0,0 | 0,0 | 0,0 | 71,0 | 36,9 |
| RPA3b | At4g18590 | 59,9 | 22,0 | 4,7 | 0,0 | 71,3 | 3,7 |
| PCNA1 | At1g07370 | 13,4 | 24,1 | 2,5 | 0,0 | 78,5 | 119,4 |
| PCNA2 | At2g29570 | 15,9 | 43,6 | 0,0 | 81,2 | 93,4 | 63,1 |
| LIG1 | AT1G08130 | 23,9 | 17,8 | 53,6 | 20,2 | 193,8 | 188,5 |
| Clamp loader |  |  |  |  |  |  |  |
| RFC1 | At5g22010 | 30,1 | 46,2 | 77,4 | 246,7 | 96,3 | 141,1 |
| RFC2 | At1g63160 | 16,6 | 31,8 | 0,0 | 0,0 | 24,8 | 22,0 |
| RFC3 | At5g27740 | 62,0 | 31,7 | 62,2 | 1,2 | 21,6 | 29,9 |
| RFC4 | At1g21690 | 39,8 | 3,8 | 26,8 | 15,1 | 30,8 | 80,2 |
| RFC5 | At1g77470 | 45,4 | 17,5 | 0,0 | 109,7 | 33,6 | 41,2 |

\* Expression data from Zhao et al. doi.org/10.1016/j.devcel.2019.04.016
\* Expression data from Susaki et al. doi.org/10.1371/journal.pbio.3001123

## Notes

### Competing Interest Statement

The authors have declared no competing interest.

